# ANI-netID: a genome similarity-network based biological identification system

**DOI:** 10.1101/2025.04.19.649685

**Authors:** S. Nozawa, K. Watanabe

## Abstract

Fungal identification is based on sequence similarity comparisons, with multiple genes often utilized in conjunction, similar to most taxonomic studies. The identification process involves the individual comparison of the similarities of each gene. However, owing to insufficient information, incomplete lineage sorting, gene transfer, hybridization, gene duplication, and loss, different species may match, depending on the DNA region used for comparison. Additionally, identification methods exist that set a uniform similarity threshold value, but they hit multiple species above the threshold. In this study, we introduced ANI-netID, which was developed to address these challenges. ANI-netID is an identification method based on the phylogenetic species concept, where the threshold value is determined by the combination of individuals with the lowest nucleotide similarity within groups (clique groups), such as species and genera. This checks whether the queried individual fall into the same clique group as the reference individual with the highest similarity. ANI-netID solves not only the above-mentioned problems but also indicates that the individual being identified may represent a new lineage, if the individual does not belong to any clique group.

## Introduction

Species identification is important in biodiversity research and exerts a significant influence on the field of applied biological research. The 1980s witnessed significant advancements in the field, with the development of methods for determining the base sequences of genes. Concurrently, several databases have been established to facilitate the storage and exchange of this information, including GenBank (National Center for Biotechnology Information base sequence database), DNA Data Bank of Japan (DDBJ), and the European Nucleotide Archive (ENA) of the European Molecular Biology Laboratory (EMBL). Currently, a substantial corpus of nucleotide sequence data has been amassed and is being used for identification purposes. In the context of research on microorganisms that possess a paucity of distinguishing characteristics, such as fungi, DNA-based identification methods have proven to be highly advantageous. These methods are frequently employed in the identification of pathogens that affect humans, animals, and agricultural crops [1].

For the identification of fungi, the Basic Local Alignment Search Tool (BLAST) selects the species name with the highest percentage of similarity.

However, the absence of a predetermined similarity threshold for selecting the species name makes it impossible to account for the potential of the identified fungus as a novel lineage. Conversely, recent methodologies have employed a threshold for identification, particularly in metabarcoding analyses. For instance, tools for species identification based on ITS sequences have been developed, such as CloVR-ITS, DAnIEL, and PIPITS [2–4].

These tools determine the species name when there are species that match the threshold value (e.g., 97% similarity) or higher for species identification. Cai and Druzhinina (2021) [5] published criteria for the identification of *Trichoderma* species based on a threshold for species identification based on similarity values. Specifically, they stated that the reference strains that match 99% or more using the second largest subunit of RNA polymerase II as an indicator or 97% or more using the translation elongation factor 1-alpha gene as an indicator, are the same species. However, some combinations demonstrated a similarity value of 99% or more between species. Consequently, the efficacy of species identification based on uniform similarity values is contingent upon the specific strain in question.

Another problem with the current species identification technique is that because species classification is currently based on multiple DNA regions in several taxa, multiple regions must also be used for species identification. This means that each region is individually analyzed using BLAST to identify species with high similarity. However, when species identification is based on multiple DNA regions, different species are often hit most similarly, depending on the region analyzed. This is due to the evolutionary history of each gene, including incomplete lineage sorting, hybridization in ancestral lineages, gene transfer, gene duplication, and loss [6]. In this case, identification cannot be based on similarity comparisons. Therefore, species must be identified by molecular phylogenetic analysis based on data combining DNA regions according to the latest taxonomic studies. First, in the context of the current taxonomy, in which multiple gene regions are combined for phylogenetic analysis, it is not surprising that discrepancies arise in the identification results obtained from the similarity analyses of individual genes.

In this study, we propose a novel identification system called the Average Nucleotide Identity Network Identification (ANI-NetID). This biological identification system is based on groups (clique groups) composed of individuals whose sequences have been annotated (e.g., groups composed of individuals of the same species). This checks whether the queried individual belongs to the same clique group as the reference individual with the highest similarity.

This system has the advantage of eliminating subjectivity because the threshold values are set according to the rules. Furthermore, it can be concluded that individuals who do not belong to any clique may belong to a new lineage. Additionally, when employing multiple gene regions in this network, the threshold is determined based on the Average Nucleotide Identity, facilitating identification. This system solves the problem of identifying different species for each gene. In this study, we demonstrated ANI-NetID through simulations using a dataset to identify the species of *Nigrospora* fungi.

## Materials and Methods

### 1. ANI-NetID pipeline

1) The reference sequence data are integrated with the query sequence for each DNA region.
2) Alignment is performed independently for each DNA region; ANI-netID uses MAFFT v7.310 [7] for multiple alignments, which creates a guide tree based on distances and aligns sequences based on this tree, which is compatible with current classification methods that build classification systems based on phylogenetic trees. Note that BLAST, which finds pairwise similarities without creating a guide tree, is not used in the next process of obtaining similarity values.
3) After alignment, the similarity values between sequences for each DNA region are determined using exhaustive pairwise sequence comparisons.
4) The average similarity of the DNA regions is obtained for all individual combinations. For example, if the homology of each region between two individuals is Locus A: 97%, Locus B: 98%, and Locus C: 99%, the average is 98.
5) The pair with the lowest similarity within each clique group is determined based on the average similarity, and its similarity value (LAC; lowest ANI value in the clique group) is obtained.
6) We then search for the reference sequence with the highest similarity to the query sequence and obtain its similarity value (HAQ: highest ANI value to the query).
7) HAQ is compared with the LAC among the combinations of individuals in the clique group to which the individual belongs and is identified based on the following **identification evaluation criteria:**

1. **HAQ =100%:** It is identified as belonging to the same clique group as an individual if it is 100% similar to a query.
2. **HAQ > LAC:** The query belongs to the clique group with the most similar individuals. In this case, the query is identified by a clique group.
3. **HAQ≦LAC:** A query is closely related to the clique group to which the most similar individuals belong, yet it does not belong to either clique group. In this case, as a phylogenetic relationship, a query may be located in the ancestral lineage of the clique group or in an independent lineage. Consequently, queries should be assessed using molecular phylogenetic analyses.

The program developed in this study (ANInetID.py; S1 Dataset) produced the following final outputs: i) the name of the reference individual most homologous to the query, ii) its similarity value (HAQ), iii) the name of the clique group to which the query belongs, and iv) the threshold value of each clique group (LAC). By truncating the results to an arbitrary value, the identification results can be visualized by depicting a network diagram in which the individuals above that value are connected. Moreover, the output of the identification process, encompassing the alignment results of the DNA regions, respective homology values, and mean values of the DNA regions utilized, can be obtained.

### 3. Experiment

Simulations were performed using the Python program (ANInetID.py; S1 Dataset) developed in this study based on a biological dataset. The taxa targeted in the simulations were *Nigrospora* spp. Five query sequences (query 1-5) were identified using sequences of 58 strains from 29 species (Table 1) that had already been identified as reference sequences. The DNA regions used as reference were the three regions the internal transcribed spacer region (ITS), transcribed elongation factor 1 alpha (*tef1*), and *β-tubulin*, which have been used in taxonomic studies of the genus. The reference sequences in each region were trimmed at both ends after multiple alignments in advance, so that the common 5‘ to 3’ end sites throughout the sequence could be identified as the reference. No individuals from the other clique groups were mixed within the clique group (S1 Fig).

**Table 1.** GenBank accession numbers of the DNA sequences used in this study.

| Species | Strains | Genebank accession no. |  |  |
| --- | --- | --- | --- | --- |
| | | ITS | β-tubulin | tef1 |
| *Apiospora malaysiana* | CBS 102053 | KF144896 | KF144988 | KF145030 |
| A.<br>*pseudoparenchymatica* | LC7234* | KY494743 | KY705211 | KY705139 |
| *Nigrospora aurantiaca* | CGMCC 3.18130* | KX986064 | KY019465 | KY019295 |
| *N. aurantiaca* | LC7034 | KX986093 | KY019598 | KY019394 |
| *N. bambusae* | CGMCC 3.18327* | KY385307 | KY385319 | KY385313 |
|  | LC7245 | KY385305 | KY385321 | KY385315 |
| *N. brasiliensis* | CMM 1214* | KY569629 | MK720816 | MK753271 |
|  | CMM 1217 | KY569630 | MK720817 | MK753272 |
| *N. camelliae-sinensis* | CGMCC 3.18125* | KX985986 | KY019460 | KY019293 |
| *N. chinensis* | LC6851 | KX986049 | KY019579 | KY019450 |
|  | CGMCC 3.18127* | KX986023 | KY019462 | KY019422 |
| *N. covidalis* | CGMCC 3.20538* | OK335209 | OK431479 | OK431485 |
|  | LC158337 | OK335210 | OK431480 | OK431486 |
|  | LC2725 | KX985960 | KY019487 | KY019313 |
|  | LC4566 | KX986022 | KY019545 | KY019354 |
| *N. endophytica* | A.R.M. 687 | OM265226 | OP572418 | OP572415 |
|  | A.R.M. 973* | OM265233 | OP572420 | OP572416 |
| *N. falsivesicularis* | CGMCC 3.19678* | MN215778 | MN329942 | MN264017 |
|  | LC13553 | MN215779 | MN329943 | MN264018 |
| *N. globospora* | CGMCC 3.20539* | OK335211 | OK431481 | OK431487 |
|  | LC15839 | OK335212 | OK431482 | OK431488 |
| *N. gorlenkoana* | CBS 480.73* | KX986048 | KY019456 | KY019420 |
| *N. guangdongensis* | CFCC:53917* | MT017509 | MT024495 | MT024493 |
| *N. guilinensis* | LC7301 | KX986063 | KY019608 | KY019404 |
|  | CGMCC 3.18124* | KX985983 | KY019459 | KY019292 |
| *N. hainanensis* | CGMCC 3.18129* | KX986091 | KY019464 | KY019415 |
|  | A.R.M.967 | OM265228 | OM793057 | OM642834 |
|  | A.R.M.968 | OM265229 | OM793058 | OM642835 |
|  | A.R.M.972 | OM265232 | OP572419 | OM642837 |
|  | A.R.M.976 | OM265236 | OM793060 | OP572417 |
| *N. lacticolonia* | CGMCC 3.18123* | KX985978 | KY019458 | KY019291 |
|  | A.R.M. 921 | OM265227 | OM642838 | OM642833 |
| *N. manihoticola* | A.R.M. 645* | OM265224 | OM869479 | OM914791 |
| *N. musae* | CBS 319.34* | KX986076 | KY019455 | KY019419 |
|  | LC6385 | KX986042 | KY019567 | KY019371 |
| *N. oryzae* | LC2724 | KX985959 | KY019486 | KY019312 |
|  | LC4265 | KX985994 | KY019518 | KY019335 |
| *N. osmanthi* | CGMCC 3.18126* | KX986010 | KY019461 | KY019421 |
|  | LC4487 | KX986017 | KY019540 | KY019438 |
| *N. pernambucoensis* | A.R.M.651 | OM265225 | OM869480 | OM914792 |
|  | A.R.M. 974* | OM265234 | OM869481 | OM914793 |
| *N.philosophiae-doctoris* | CGMCC 3.20540* | OK335214 | OK431484 | OK431490 |
| *N. pyriformis* | CGMCC 3.18122* | KX985940 | KY019457 | KY019290 |
|  | A.R.M.970 | OM265231 | OM642839 | OM513904 |
| *N. rubi* | LC2698* | KX985948 | KY019475 | KY019302 |
| *N. saccharicola* | LC12057 | MN215789 | MN329952 | MN264028 |
|  | CGMCC 3.19362* | MN215788 | MN329951 | MN264027 |
| *N. sacchari-officinarum* | CGMCC 3.19335* | MN215791 | MN329954 | MN264030 |
|  | LC13531 | MN215792 | MN329955 | MN264031 |
| *N. singularis* | CGMCC 3.19334* | MN215793 | MN329956 | MN264032 |
|  | LC12068 | MN215794 | MN329957 | MN264033 |
| *N. sphaerica* | LC2839 | KX985964 | KY019491 | KY019317 |
|  | LC2840 | KX985965 | KY019492 | KY019318 |
| *N. vesicularis* | LC0322 | KX985939 | KY019467 | KY019296 |
|  | CGMCC 3.18128* | KX986088 | KY019463 | KY019294 |
| *N. vesicularifera* | CGMCC 3.19333* | MN215812 | MN329975 | MN264051 |
|  | A.R.M.975 | OM265235 | OM642840 | OM513905 |
| *N. zimmermanii* | CBS 290.62* | KY385309 | KY385317 | KY385311 |
|  | CBS 984.69 | KY385310 | KY385322 | KY385316 |

ITS = internal transcribed spacers; *tef1* = translation elongation factor 1-alpha gene.
*Apiospora malaysiana* and *A. pseudoparenchymatica* were used for only phylogenetic
analysis as out group. Ex-type strains are marked with an asterisk. All the query sequences
are available in S1 dataset.

**Table 2.**
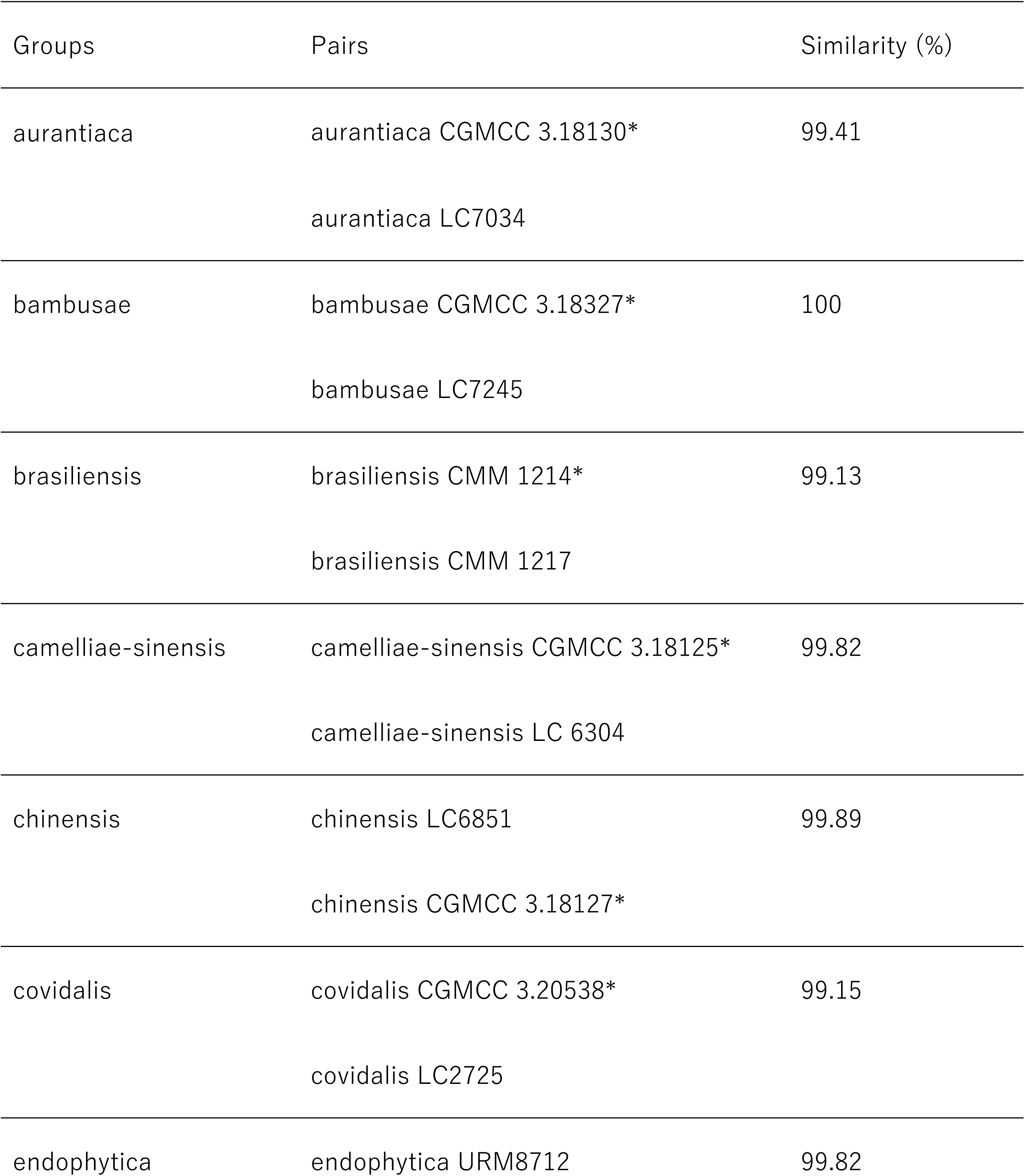
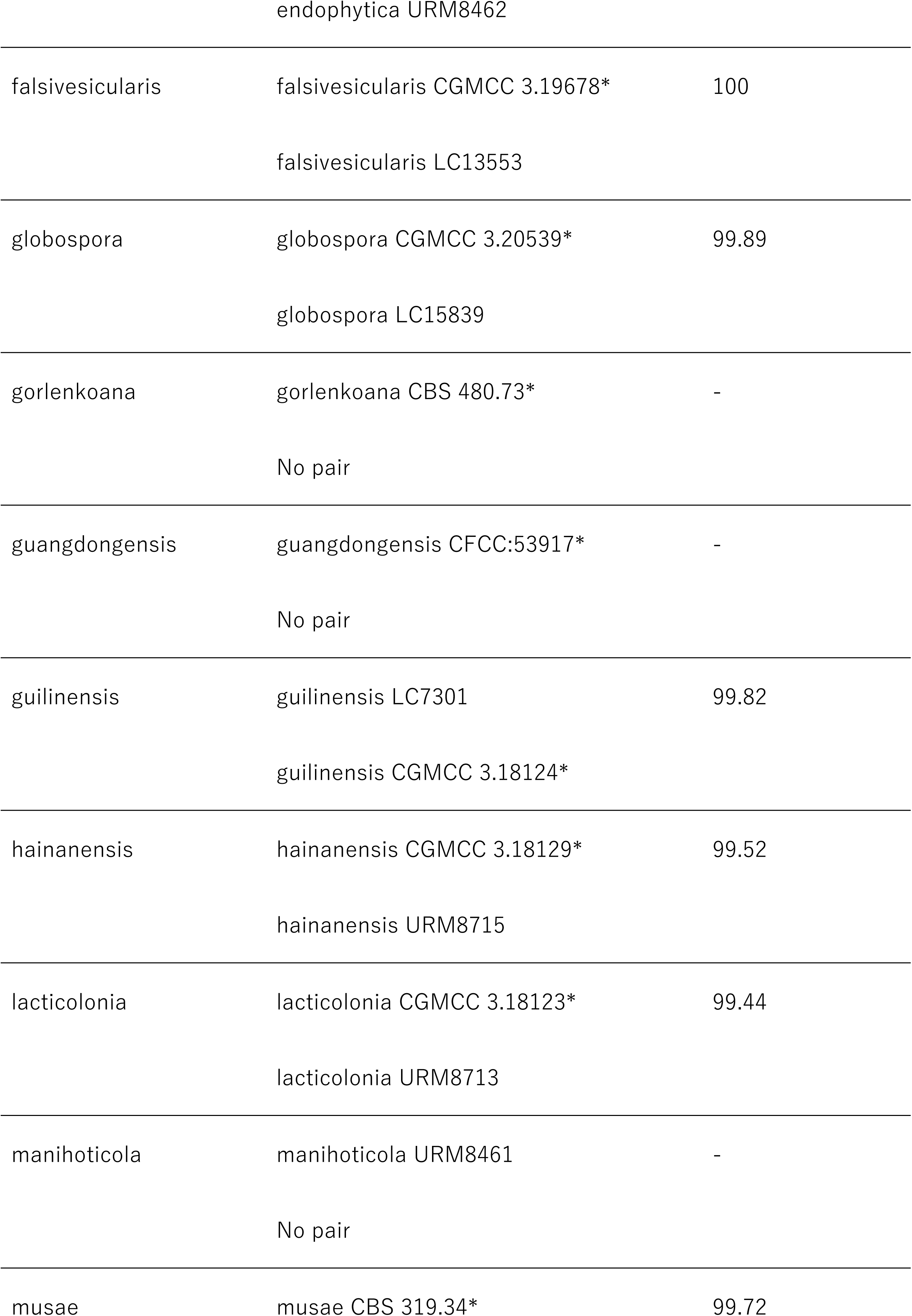
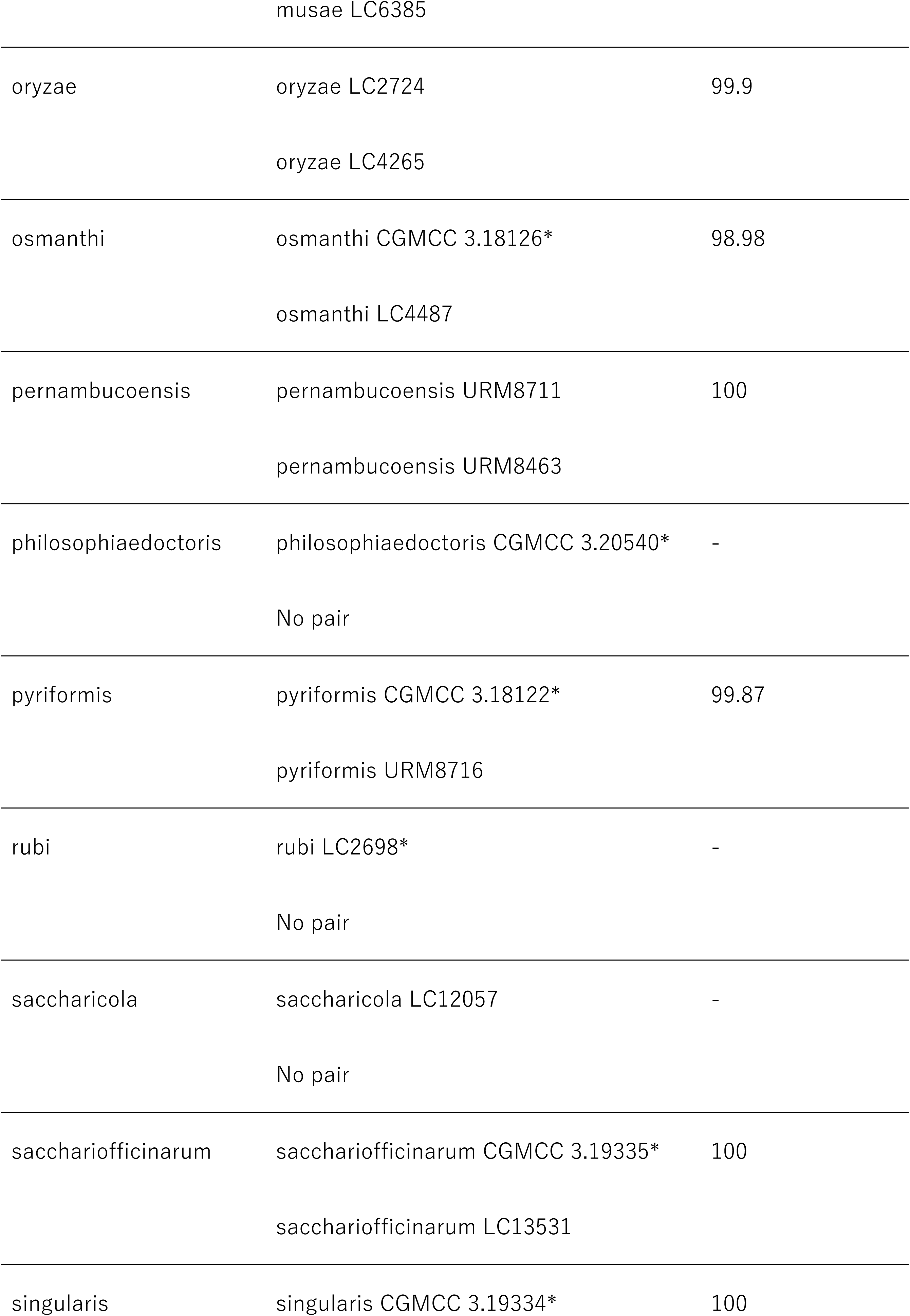
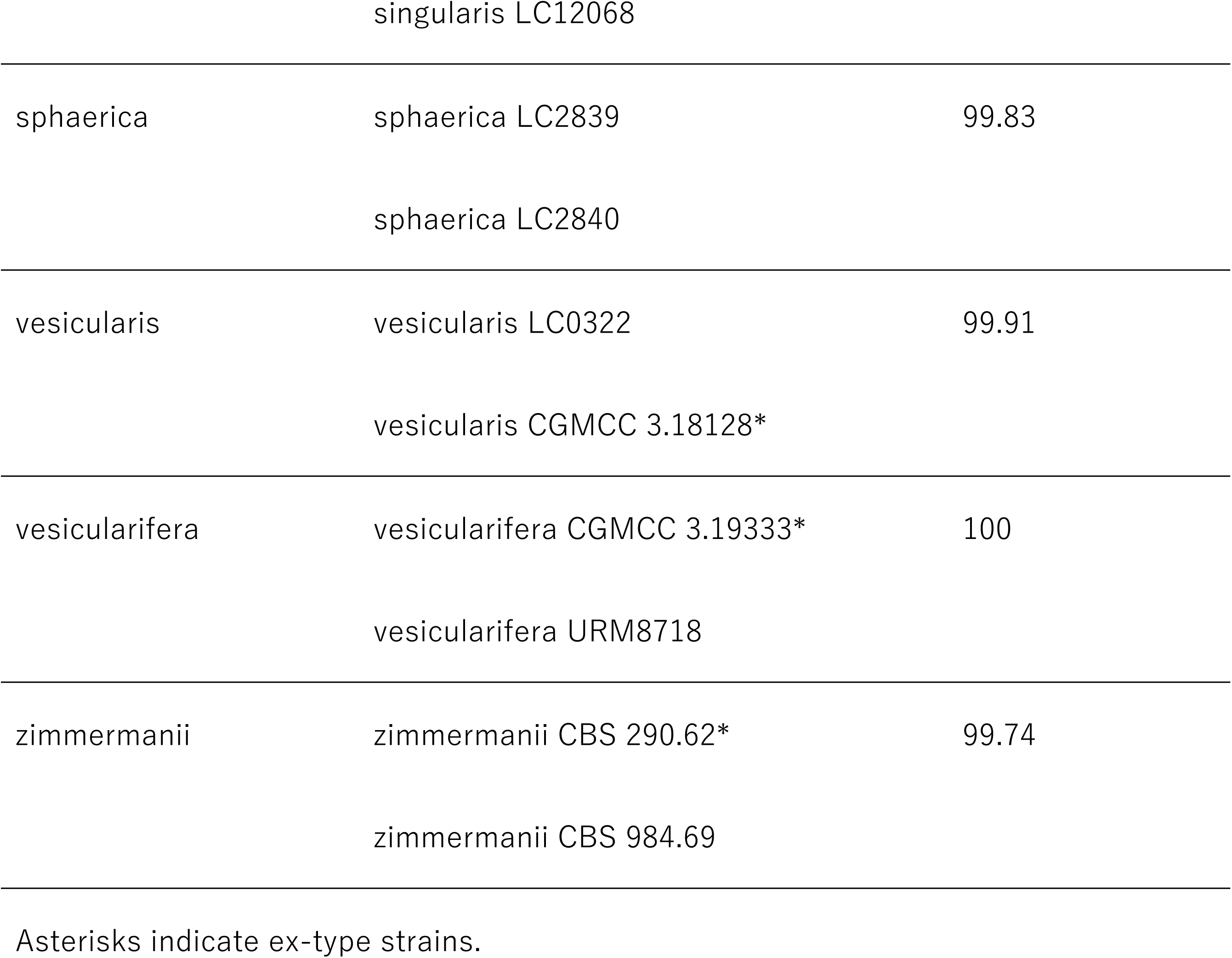
Pairs with lowest identities in each clique group (species) and their identity.

#### Phylogenetic analyses

Phylogenetic analyses were performed to evaluate the identification results obtained using ANI-NetID. To construct phylogenetic trees, three query strains and 59 strains of *Negrospora* spp., including ex-type strains, which were from the same dataset as that for ANI-NetID, were used (Table 1). As out groups, *Apiospora malaysiana* and *A. pseudoparenchymatica* were used. All the ITS, *β-tubulin*, and translation elongation factor 1-alpha gene (*tef1*) sequences were used for multiple alignments by MAFFT v7.310 and were concatenated into a dataset. Three phylogenetic analyses—neighbor-joining (NJ), maximum-likelihood (ML), and maximum-parsimony (MP) methods—were performed using MEGA 10 [8]. The reliability of each internal branch in the phylogenetic trees was tested using bootstrap analysis [9] with 1,000 random addition replicates. All sites with gaps were treated as having missing data. For the ML tree, the best substitution model was selected based on the Akaike information criterion in MEGA 10.

## RESULTS

### Identification of query 1

The reference strain with the highest similarity to query 1 was *N. lacticolonia* strain 001, with a value of 99.91% (Fig 2a; Link HAQ). This value exceeded the similarity value of the least similar combination within the *N. lacticolonia* clique group, CGMCC 3.18123 and URM8713, namely 99.44%; Fig 2a; Link LAC). Thus, the queried strain was identified as being *N. lacticolonia*.

**Fig 1.**
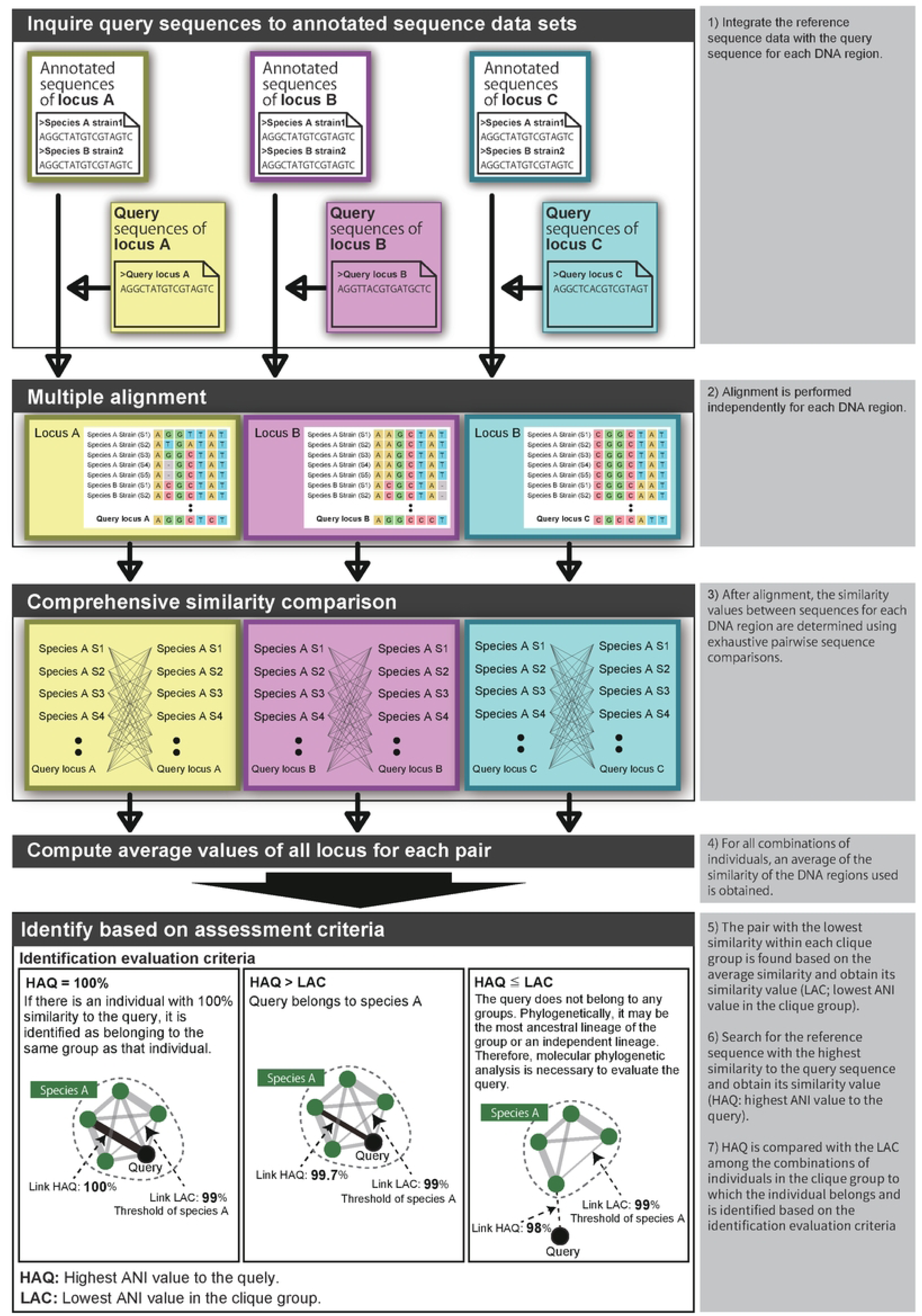
Pipeline of ANI-netID.

**Fig 2.**
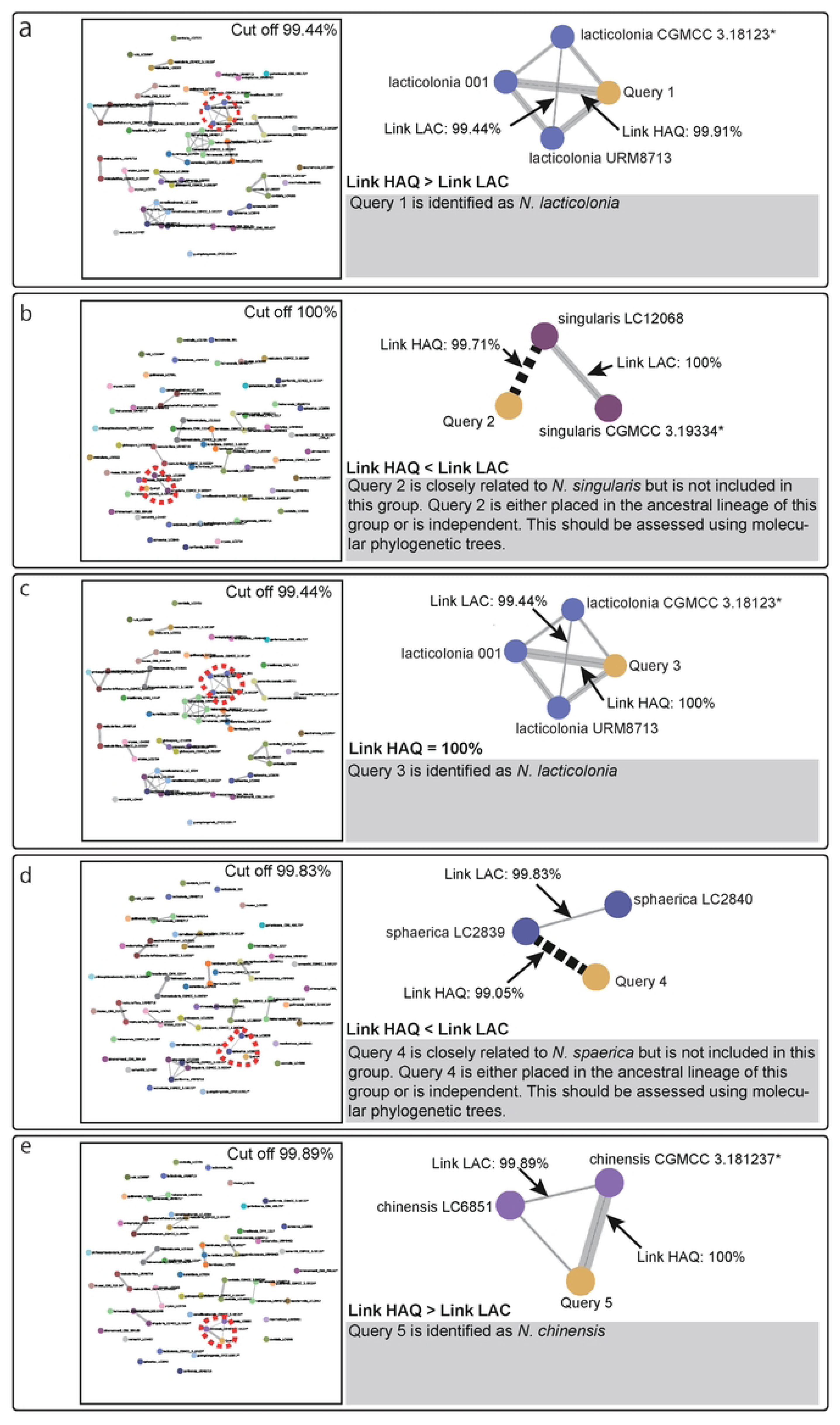
Results of identification based on ANI-netID. a) Query 1, b) Query 2, c) Query 3, d) Query 4, e) Query 5. Connected individuals have a higher value than the cut-off value. The dashed line connects the query to the individual with the highest similarity, indicating that the query does not belong to the same group as that individual. LAC: lowest ANI value in the clique-group, HAQ: highest ANI value to the query.

### Identification of query 2

Query 2 identification showed that this strain had the highest similarity (99.71 %) to *N. singularis* LC12068 (Fig 2b; Link HAQ). This value was less than the clique group threshold for *N. singularis* with 100% similarity between *N. singularis* LC12068 and CGMCC 3.19334 (Fig 2b; Link LAC). Therefore, query 2 does not belong to *N. singularis* but is closely related to *N. singularis*. Consequently, it is plausible that this lineage represents the most ancestral lineage of *N. singularis*. Alternatively, it may be a novel independent lineage. Consequently, the evaluation of query 2 by molecular phylogenetic and morphological analyses is necessary to determine its taxonomic position.

### Identification of query 3

The reference strain with the highest similarity to query 3 was *N. lacticolonia* 001 with a value of 100% (Fig 2c; link HAQ). Therefore, query 3 was identified as *N. lacticolonia*.

### Identification of query 4

Query 4 identification demonstrated that the reference strain to which this strain is most similar is *N. sphaerica* LC2839, with a value of 99.05% (Fig 2d; Link HAQ). This value is below the *N. sphaerica* clique-group threshold of 99.83% similarity between *N. sphaerica* LC2839 and LC2840 (Fig 2d; Link LAC). Consequently, query 4 does not belong to *N. sphaerica* but is a closely related lineage. Consequently, query 4 may represent either the most ancestral lineage of *N. sphaerica* or a new, independent lineage. Consequently, the evaluation of query 4 by molecular phylogenetic and morphological analyses is required.

### Identification of query 5

The reference strain with the highest homology to query 5 was *N. chinensis* CGMCC 3.18127, with a value of 100% (Fig 2e; Link HAQ). Thus, the query strain was identified to be *N. chinensis*.

### Evaluation of the identification result

To evaluate the identification results from ANI-NetID, molecular phylogenetic analysis was performed on the dataset that had been used for identification and the five query strains, as well as the outgroups *Apiospora malaysiana* and *A. pseudoparenchymatica*, based on combined ITS, *β-tubulin*, and *tef1* data (Table 1). Following alignment, 1,148 sites were included, comprising 515 variable sites, 445 parsimony-informative sites, and 69 singleton sites. In the resulting ML, MP, and NJ phylogenetic trees, the strains of the same species (clique group) formed a monophyletic lineage (Fig 3).

**Fig 3.**
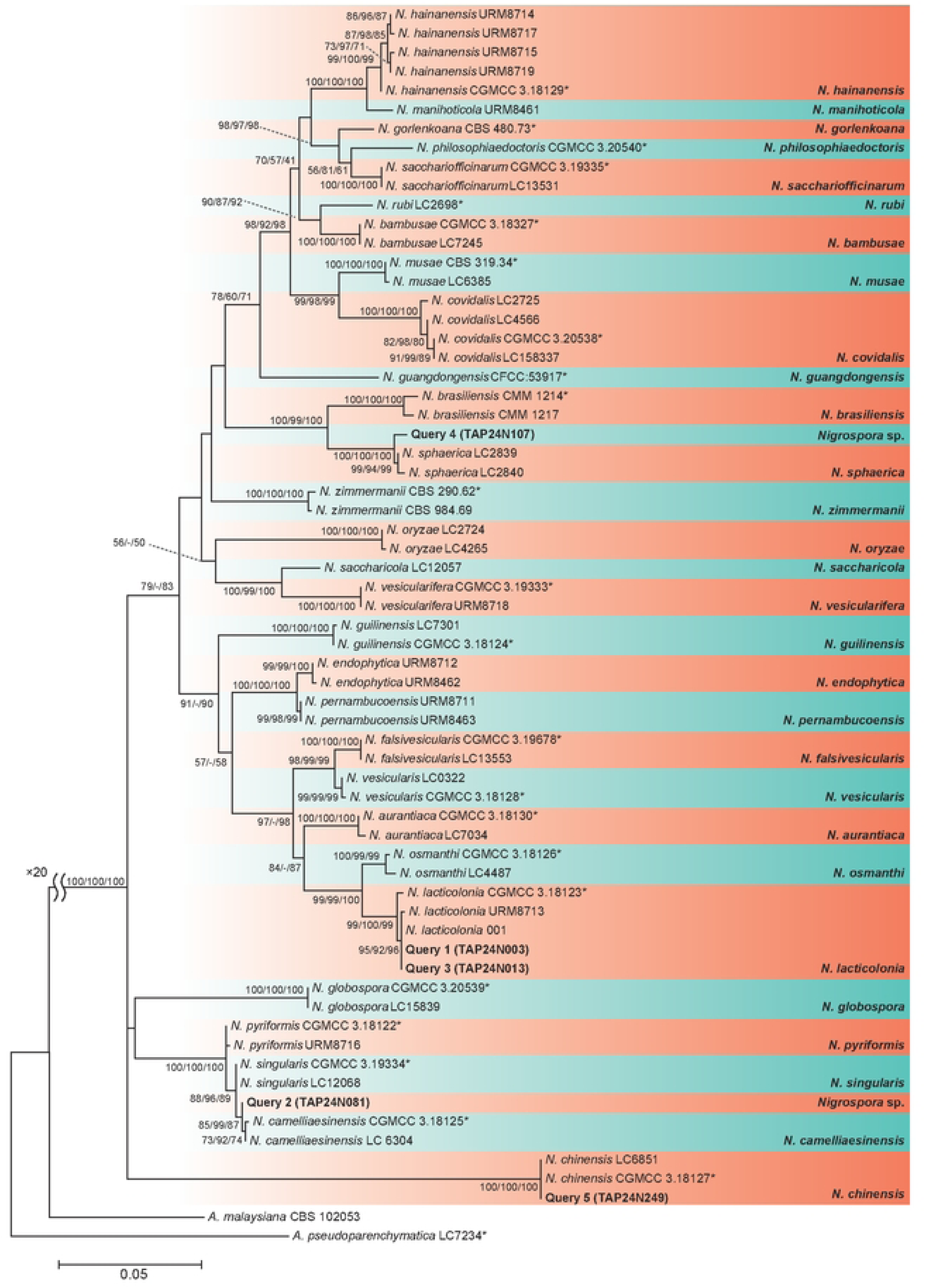
Maximum-likelihood phylograms of *Nigrospora* species inferred from the concatenation of the ITS, *β-tubulin*, and *tef1* genes. Values on the branches are ML, MP, and NJ bootstrap values (ML/MP/NJ). Strains with an asterisk are Ex-type strains. Queries are highlighted in bold.

Queries 1 and 3 belonged to the *N. lacticolonia* clade (Fig 3; bootstrap values BS, ML/MP/NJ: 97/99/100). Within this clade, both the strains formed a monophyletic group with *N. lacticolonia* 001 (Fig 4a). Query 2 was a sister to *N. camelliae-sinensis* and shared a common ancestor with *N. singuralis* (Fig 3; ML/MP/NJ: 88/96/89; Fig 4b). Query 4 was the ancestral lineage of the *N. sphaerica* clade (Fig 3; ML/MP/NJ: 100/100/100; Fig 4c). Query 5 belonged to the *N. chinensis* clade (Fig 3; ML/MP/NJ: 99/100/100; Fig 4d). These results support the ANI-netID results (Fig 2).

**Fig 4.**
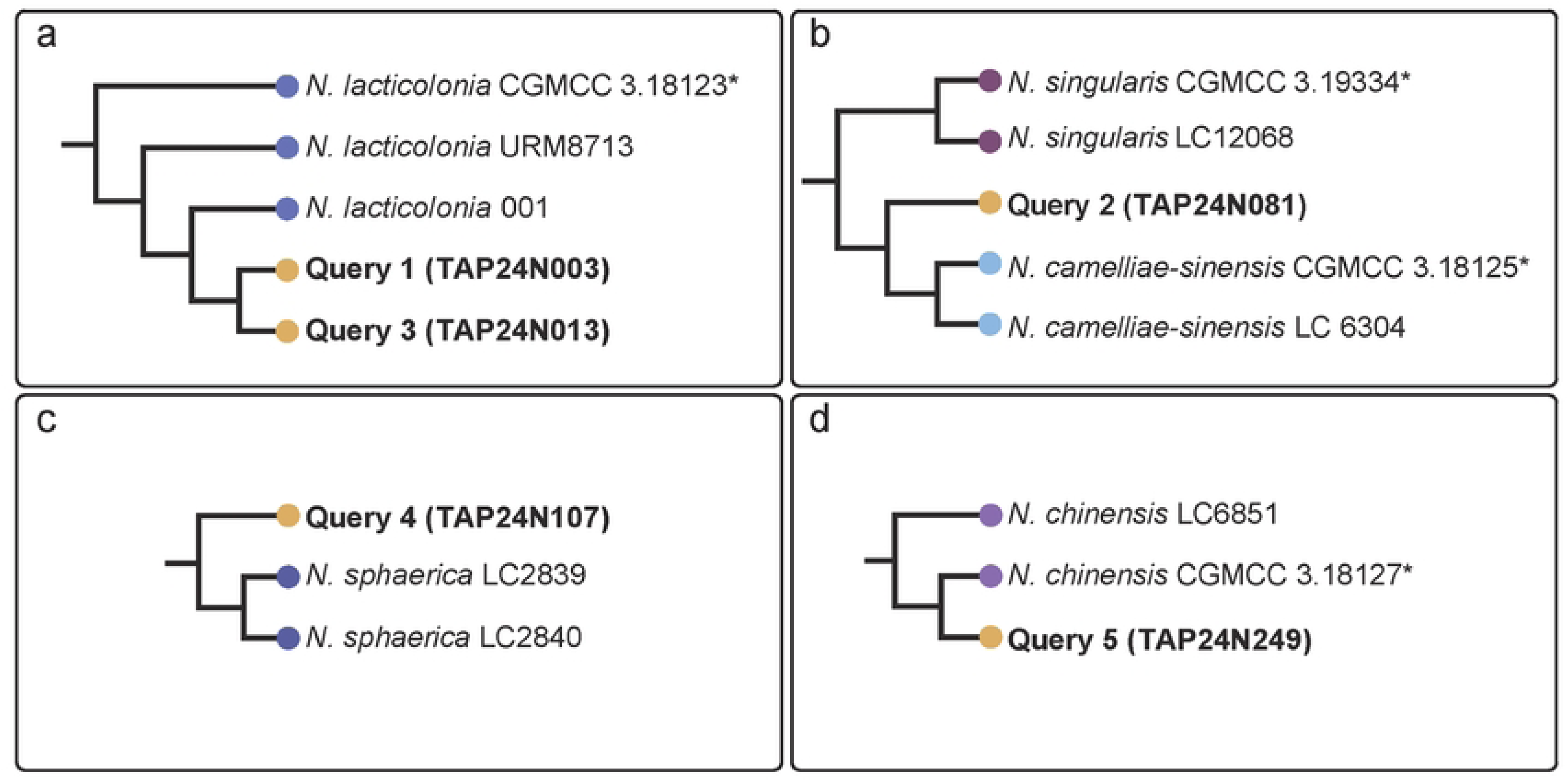
Parts of the maximum-likelihood cladograms (see Fig 3). *N. lacticolonia* clade (a), *N. camelliae-sinensis* and *N. singularis* clades (b), *N. sphaerica* clade (c), and *N. chinensis* clade (d). Strains with an asterisk are ex-type strains. Queries are highlighted in bold. The branch length is not related to evolutionary distance.

## Discussion

This study introduces a new identification system called ANI-netID. This method differs from conventional identification methods in that it is based on a “clique-group” unit. This is a group of individuals of a species or genus based on similarity values between individuals. In this system, the similarity threshold for identification is automatically determined for each clique group. In other words, the similarity value between the two least similar individuals in the clique group is the threshold. This clarifies whether the individual to be identified belongs to the clique group. This addresses the issue of species identification failure caused by multiple high-scoring hits in BLAST searches [5]. In addition, the system uses average similarity when multiple DNA regions are used. This overcomes the problem of BLAST searches in individual regions for species identification, resulting in hits for different species in different regions.

However, as with the DNA barcoding method for species identification, there is no universal DNA region common to all life forms [10]. Therefore, this system requires identification of the DNA regions used for each taxon. Therefore, the pre-determination of higher taxonomic ranks should be based on morphological analysis or the ITS region.

In this study, an ANI-netID system was constructed for *Nigrospora* spp. and species identification was attempted for five strains based on their ITS, *TUB* and *TEF* regions. The identification results of this system were consistent with the results of the molecular phylogenetic analysis (Figs 2, 3, and 4). The system uses the rate of matching nucleotide sites between two sequences as its basis. This has the same meaning as the p-distance, which calculates the rate of different nucleotide sites. The distances between individuals were similar to the phylogenetic relationships based on the p distance. Therefore, this system is compatible with the taxonomic systems established by molecular phylogenetic analysis. This likely produced results reflecting the taxonomic system.

However, biases in nucleotide substitution rates due to codon positions [11–16], etc. are not considered. Therefore, the ANI-netID identification system cannot be used in cases where the grouping of taxa constructed by phylogenetic analysis based on complex mathematical models is different from that obtained by the p distance (e.g., when the same species is polyphyletic). Developers must ensure that the groupings for each target taxon match the groupings of the system before building the system. As a first step, it is recommended to check whether the phylogenetic tree based on the p distance is consistent with the current classification. The program developed in this study included a simple tool to check groupings (S1 dataset). As shown in Fig 1 S, it can be verified that populations that should belong to the same clique-group can be grouped without contamination of individuals from other clique-groups by exceeding the threshold.

Although a simulation of species identification was conducted in this study using *Nigrospora* sp. as an example, the development of this identification system cannot be applied only to species identification. For example, three genera related to Pestalotiopsis (*Pestalotiopsis*, *Neopestalotiopsis* and *Pseudopestalotiopsis*) are similar in their branchial morphology [17], making them difficult to determine. In such cases, a system can be created to determine the genus based on the ANI-netIDs. In addition, agricultural and clinically important fungi such as the genera *Colletotrichum* and *Fusarium* are taxonomically studied intensively and have a large number of species. Species identification is conducted in molecular phylogenetic groups called species complexes, which are established under the genus [18, 19]. Therefore, it is necessary to determine the species complexes before species identification. This system is expected to be effective in such cases.

In addition to ANI-netID, an important aspect of species identification is the accumulation of longer sequences as data. The longer the sequence, the more information (mutations) that can be used to identify the individual and the more accurate the identification [20]. Recent developments in next-generation sequencing technologies have made it possible to obtain sequences that are longer than traditional barcode regions. Taxonomic studies have also begun to consider taxonomic systems based on phylogenetic relationships derived from genome-scale analyses [21, 22]. In the future, as more genomic data accumulate, more taxa will be organized. This will increase the amount of data available for ANI-netID, and longer sequences will become available, leading to improved identification accuracy.

We plan to make ANI-netID a platform where any taxonomist can develop a system for each taxonomic group individually and provide access to users who need identification work. This will allow more data to be accumulated and identified using ANI-netID for various taxonomic groups.

## Supporting Information

**S1 Fig.** Networks formed by eliminating the threshold values.

**S1 Dataset**. Programs and tested datasets.

## Acknowledgments

We thank Dr. Yosuke Seto of the Cancer Chemotherapy Center, Japanese Foundation for Cancer Research, and Dr. Takayuki Aoki of National Agriculture and Food Research Organization for their invaluable advice.

## Author Contributions

**Conceptualization:** Kyoko Watanabe, Shunsuke Nozawa.

**Formal analysis:** Shunsuke Nozawa.

**Funding acquisition:** Kyoko Watanabe.

**Investigation:** Kyoko Watanabe, Shunsuke Nozawa.

**Methodology:** Shunsuke Nozawa.

**Project administration:** Kyoko Watanabe.

**Software:** Shunsuke Nozawa.

**Supervision:** Kyoko Watanabe.

**Validation:** Kyoko Watanabe, Shunsuke Nozawa.

**Visualization:** Shunsuke Nozawa.

**Writing – original draft:** Shunsuke Nozawa.

**Writing – review & editing:** Kyoko Watanabe.

## Notes

### Competing Interest Statement

The authors have declared no competing interest.

